# Easy Removal of Steric Clashes and Entanglements in Macromolecular Systems by Temporary Addition of a Fourth Spatial Dimension

**DOI:** 10.1101/2023.04.26.537866

**Authors:** Adrian H. Elcock

## Abstract

When models of complicated macromolecular systems are constructed, it is common to inadvertently include either gross steric clashes or entanglements of extended loop regions. Removing these problems with conventional energy minimization or dynamics algorithms can often be difficult. Here I show that one easy alternative is to temporarily add an extra spatial dimension and to displace atoms or molecules along this fourth dimension such that the distances between atoms, when measured in 4D, are no longer considered clashing. Adding in half-harmonic potential functions to mimic walls in this 4^th^ dimension, and then moving these walls toward each other, has the effect of decreasing the space available in the 4^th^ dimension and drives atoms to avoid each other in 3D. I illustrate the method with three examples: two showing how interlocked ring polymers can be easily disentangled from each other in both 2D and 3D, and one showing how ten identical coarse-grained protein models, all placed at the same point in 3D space, can be separated from each other, without distorting their structures, during the course of a single energy minimization. A sample program implementing the method is available that can be easily adapted to other situations.

## Introduction

In recent years, there has been an increase in efforts to build detailed structural models of very large macromolecular systems such as those encountered in biological cells. These range from complicated protein assemblies, such as the nuclear pore complex [1] and bacterial chemoreceptor arrays [2], to large RNAs containing a few thousand nucleotides [3], to models of entire chromosomes, at least one bacterial version of which has been modeled at single-nucleotide resolution [4]. One common issue that is encountered when building detailed models of large molecules that contain extended flexible elements – especially complicated RNAs and DNAs – is that regions of the structures that are non-local in sequence can become entangled with each other in ways that are of questionable realism. This tendency has already been noted as a common feature of predicted RNA models by the Szachniuk group [5], who have developed methods to automatically identify (but not resolve) such entanglements, and our experiences with attempting to build models of large RNAs using existing methods have produced similar results [3]. In the case of RNAs, many of the entanglements are both obvious and clearly unrealistic: e.g. single strands of RNA can sometimes be found poking through the middle of double-helical regions in ways that would not be tolerated in a real structure. In other cases, however, entanglements can be more subtle, and the extent to which they are unrealistic may be a matter of debate.

A related issue that often arises with the construction of large, complicated molecular models is the presence of steric clashes that can be difficult or impossible to resolve. Depending on the situation, there are at least two ways that such clashes could, in principle, be removed. One possible approach that would likely be effective for systems that consist of free macromolecules that are primarily globular in shape would be to seek to remove steric clashes by: (1) temporarily shrinking all molecules (e.g. by scaling down all of their bond lengths), (2) repositioning them such that the shrunken versions of the molecules are all free of clashes, and (3) running a series of energy minimizations over the course of which bond lengths are gradually restored to their original values. Such an approach is, however, unlikely to be effective when the steric clashes or entanglements in question are intramolecular in origin (e.g. as might be found in whole chromosome models) since, in such cases, no amount of shrinking a molecule will relieve it of its clashes or entanglements. In addition, such an approach requires that the user has the ability to reconstruct an initial configuration of their system “from scratch”. In many cases, however, the user may not have this freedom and may instead be faced with a situation of removing steric clashes from a pre-made system (e.g. perhaps one downloaded from elsewhere, or built by someone else). A second way by which clashes and/or entanglements could potentially be resolved is to temporarily reduce their severity by decreasing the effective radii of the clashing atoms (or, more generally, “beads”), and again running a series of energy minimizations over the course of which the radii are gradually returned to their original values. This approach has the potential to be effective in certain situations, but unless some attempt is made to ensure that the previously clashing elements move apart from each other (e.g. by applying an additional potential function to their centers of mass), there remains a significant probability that they will remain entangled (or will become entangled) with each other during the process. Moreover, when the clashing elements are non-spherical in shape, potential functions additional to a center-of-mass restraint would likely be required to ensure that all parts of the structures are relieved of steric clashes.

While it is certainly possible to imagine obtaining better results by combining the two above approaches, I describe here a simpler method for removing both entanglements and egregious steric clashes that requires minimal intervention from the user and that requires no *ad hoc* adjustment of either bond lengths or bead radii. Importantly, the approach is easy to implement since it requires only the temporary addition of an extra spatial dimension to existing molecular modeling codes. The structure of this manuscript is as follows. The Methods section outlines the structural and energetic details of the three model systems that are used to illustrate the approach. The Results section describes the conceptual basis of the approach and illustrates how it allows entanglements to be resolved in 2D and then 3D, before showing how it also allows atrocious steric clashes to be easily removed. Finally, the Discussion section suggests some possible areas of application and future development for the method.

## Methods

### Systems considered

The present work considers two simple molecular systems. The first is a pair of interlocked ring-shaped molecules, each of which consists of a ring of 24 beads connected by bonds (see below). We illustrate the method proposed here by first seeking to disentangle these two molecules in 2D, which requires temporarily adding a 3^rd^ dimension, and then in 3D, which requires temporarily adding a 4^th^ dimension (see Results). This system, therefore, serves as a simple case-study likely to be applicable to more complex molecules (such as large RNAs or DNAs) for which entanglements of extended, closed elements are common. The second system studied is a heavily clashing arrangement consisting of ten coarse-grained copies of a 24-bead protein, all placed directly on top of each other in the same region of 3D space. This system, therefore, provides a demonstration of the method’s ability to resolve steric clashes that might be accidentally built in to large-scale, crowded systems during their construction.

### Energetic models

Throughout this work we consider only simple energetic models of the kinds that are typically used in molecular mechanics codes [6, 7]. Bonds between beads are described using harmonic potential functions of the form E_bond_ = 0.5 k_b_(r – r_eq_)^2^ where E_bond_ is the energy of the bond, k_b_ is the force constant, r_eq_ is the bond’s length at equilibrium, and r is the current distance between the two beads. For the ring molecules studied here, angle terms are also added to help maintain their circular shapes during the energy minimizations. Angle terms are applied to all pairs of bonds that share a common bead and are described using harmonic potential functions of the form E_angle_ = 0.5 k_a_ (θ – θ _eq_)^2^ where E_angle_ is the energy of the angle, k_a_ is the force constant, θ _eq_ is the bond angle at equilibrium, and θ is the current angle. All bond and angle terms are calculated in the original spatial dimensions specified for each system: they are not calculated in the additional spatial dimension.

In all of the energy minimizations discussed here, interactions between bead pairs that are not connected by bonds or angles are modeled using a purely repulsive steric potential function of the form E_steric_ = ε σ_bead_^12^ / r^12^ where E_steric_ is the energy of the interaction, ε is an energy parameter, σ_bead_ is the effective bead diameter, and r is the current distance between the two beads. Importantly, all steric interactions are calculated in the space defined by the additional spatial dimension: for the systems that make use of a temporary 4^th^ spatial dimension, for example, the inter-bead distance used to determine the steric interaction is measured as the square root of Δx^2^ + Δy^2^ + Δz^2^ + Δw^2^, where w represents the 4^th^ spatial dimension.

To confine beads within a sub-region of the space defined by the extra spatial dimension, so-called “wall” functions are added with the basic form E_wall_ = 0.5 k_wall_ (u – u_ref_)^2^, where E_wall_ is the energy of the bead-wall interaction, k_wall_ is a force constant, u_ref_ is the reference position of the wall in the added spatial dimension, and u is the current position of the bead in that dimension. Two wall functions are applied. To prevent beads moving to coordinates that are more positive than the reference position +u_ref_, a wall function is applied such that E = E_wall_ when u > +u_ref_, and E_wall_ = 0, when u ≤ +u_ref_. To prevent beads moving to coordinates that are more negative than the reference position –u_ref_, a wall function is applied such that E = E_wall_ when u < –u_ref_, and E_wall_ = 0, when u ≥ –u_ref_. To relieve entanglements and steric interactions in a 2D (x,y) coordinate space, the wall potential functions act along the temporary z-coordinate; to relieve entanglements and steric interactions in a 3D (x,y,z) coordinate space, the wall potential functions act along the temporary w-coordinate. Finally, to force atoms to return to a space without the added spatial dimension, the reference positions of the two wall functions are gradually moved toward each other during simulation: when the two reference positions finally coincide, the bead coordinates in the added dimension are all set to zero.

### Displacing molecules in the temporary spatial dimension

A key initial step in the methodology developed here is to ensure that molecules are displaced sufficiently in the added spatial dimension that they are no longer entangled with each other and/or no longer experience steric clashes. There are several ways by which this might be achieved. In the present case, which considers only simple systems, I have chosen to apply displacements that are known to be sufficient to ensure that all bead-bead distances for all nonbonded interactions exceed σ_bead_ (see below). In more complicated cases – such as those that we have recently encountered when applying the methodology to large RNAs [3] – we have found it effective to apply repeated random displacements to each molecule in the added spatial dimension until the bead-bead distances for all nonbonded interactions exceed σ_bead_. An important point to note in doing this is that the same displacement is applied to all beads of a given molecule or structural unit: this means that, in effect, each such unit is subject to a rigid body translation in the temporary spatial dimension.

### Separation of interlocked rings in 2D by temporary addition of a 3^rd^ dimension

To illustrate the concept underlying the method presented here, the first system examined contains two ring-shaped molecules interlocked in 2D. Each molecule contains 24 beads, separated from their nearest neighbors by 4 Å. Both the radius of each ring molecule and the initial separation of the centers of their rings are set to 15.28 Å which means that the perimeter of each ring passes through the center of the other ring (see Figure 1). Bonds were added between all nearest neighbor beads (with r_eq_and k_b_set to 4 Å and 10 kcal/mol/Å^2^, respectively), and angles were added for all pairs of bonds that shared a common bead (with θ_eq_and k_a_set to 2.880 rad (i.e. 165°) and 10 kcal/mol/rad^2^, respectively). For beads in different molecules, steric interactions were calculated with σ_bead_= 20 Å and ε = 10 kcal/mol, but were truncated (i.e. set to zero) beyond a distance of 50 Å. No intramolecular steric interactions were calculated.

**Figure 1.**
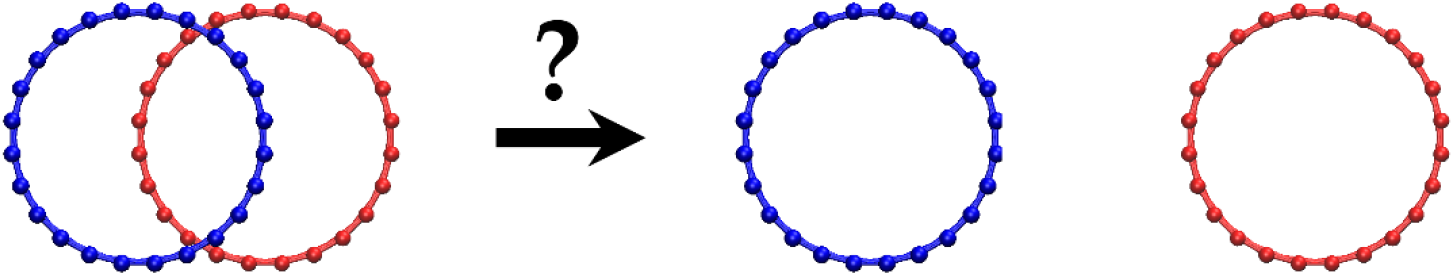
Statement of the problem for a 2D system. The goal is to separate the two interlocked ring molecules such that they remain in 2D but are no longer entangled.

While the two ring molecules are interlocked in 2D, they can be disentangled from each other if they are displaced relative to each other in a 3^rd^ spatial dimension (see Results). To accomplish this and thereby generate a suitable initial configuration of the system, all atoms in molecule 1 were assigned an initial z-coordinate of +25 Å, and all atoms in molecule in 2 were assigned an initial z-coordinate of -25 Å. Two half-harmonic wall functions were added along the z-coordinate with reference positions initially placed at z_ref_ = +25 Å and –25 Å, respectively. The force constants for these wall functions, k_wall_, were both set to 1000 kcal/mol/Å^2^. An energy minimization of the system was then carried out for 11 million steps, with the reference positions of both walls being displaced in increments of 2.5 × 10^−6^ Å at each of the first 10 million steps until they reached z_ref_ = 0. For the last 1 million steps of the energy minimization, the 3^rd^ dimension was eliminated and all steric calculations were performed in 2D. This energy minimization were carried out using Fortran code written by the author (see Code Availability).

### Separation of interlocked rings in 3D by temporary addition of a 4^th^ dimension

The next system examined used the same two two ring-shaped molecules, but starting in an initial configuration in which they are interlocked in 3D. In this case, the two ring molecules can be disentangled from each other if they are displaced relative to each other in a 4^th^ spatial dimension (see Results). To accomplish this and thereby generate a suitable initial configuration of the system, all atoms in molecule 1 were assigned an initial w-coordinate of +25 Å, and all atoms in molecule in 2 were assigned an initial w-coordinate of −25 Å. Two half-harmonic wall functions were added along the w-coordinate with reference positions initially placed at w_ref_ = +25 Å and −25 Å, respectively. As before, an energy minimization of the system was then carried out to drive the two molecules to return to 3D. All other parameters and settings for this energy minimization were identical to those of the 2D system.

### Relief of steric interactions in a clashing protein system by temporary addition of a 4^th^ dimension

The final system examined contained ten copies of a CG protein model directly superimposed in 3D. The protein is the 149-residue SFVP (Semliki Forest Virus Protein), for which we have developed a variety of CG models in a separate study [8]. The CG model used here contained 24 beads, with each bead bonded to at least four other beads. All bond force constants were set to 10 kcal/mol/Å^2^ and angle terms were omitted since the dense network of bonds was sufficient to maintain the shapes of the proteins during energy minimization. For beads in different molecules, steric interactions were calculated with σ_bead_ = 20 Å and ε = 10 kcal/mol, but were truncated beyond a distance of 50 Å; no intramolecular steric interactions were calculated. To generate a non-clashing initial configuration of the system, each molecule was subjected to a displacement in the 4^th^ spatial dimension sufficient to ensure that all molecules were separated from each other by at least 50 Å. To achieve this, molecules 1 through 10 were assigned initial w-coordinates of −225, −175, −125, −75, −25, +25, +75, +125, +175, and +225 Å, respectively. Two half-harmonic wall functions were added along the w-coordinate with reference positions initially placed at w_ref_ = +250 Å and –250 Å, respectively; as before, the force constants for these wall functions, k_wall_, were both set to 1000 kcal/mol/Å^2^. All other parameters and settings for this energy minimization were identical to those of the 4D energy minimization of the two ring molecules (see above).

### Code availability

All computer code necessary to run the energy minimizations described here will be made available to reviewers at the time of manuscript review. Upon acceptance of the manuscript for publication, the computer code will be available to the community at the following GitHub repository (https://github.com/Elcock-Lab/4D_disentangle).

## Results

The approach developed here involves solving entanglement and steric clash problems in 3D space by temporarily adding a 4^th^ spatial dimension. Since events occurring in 4D space can be difficult to visualize, a more straightforward way to illustrate the conceptual basis of the method proposed here is to instead consider how entanglement or steric clash problems in 2D space can be solved by temporarily adding a 3^rd^ spatial dimension: once this is understood, the extension to resolving similar problems in 3D space follows naturally.

### Illustration of the principle

An example system illustrating the basic problem considered here is shown in Figure 1. Here we see two ring molecules, each consisting of 24 “beads” connected by bonds to which harmonic potential functions are applied. The two rings are interlocked (entangled), and are restricted to a 2D space (the x,y plane), such that the rings are coplanar. We wish to achieve two goals. The first is to develop a method that can resolve (i.e. remove) the entanglement of the two rings using some kind of energy minimization that can be implemented in a conventional molecular mechanics code. The second is to design the method so that it is effectively automatic and therefore requires minimal intervention by the user; we make this a requirement since the areas where we anticipate applying the method are likely to be much more complex than the example system shown.

The conceptual path that leads to such an algorithm can be illustrated as follows. If we imagine performing a conventional energy minimization of the 2D system shown in Figure 1 with steric interactions and harmonic bond potentials both present, it should be clear that no disentanglement of the two ring molecules could occur. This would remain true even if we used molecular dynamics simulation instead: any attempt of the two molecules to move apart from each other would be resisted by unavoidable steric interactions between their beads. But now let us consider what would happen if we were to: (a) temporarily add a 3^rd^ spatial dimension (z), and (b) displace one or both of the two molecules along the z-axis until all beads of the two molecules are well separated from each other according to distance measurements carried out in 3D. Importantly, this can be achieved by translating all atoms of the moved molecule by an *identical* distance in z. Now the system will appear as shown on the extreme left hand side of Figure 2, from which it can be seen that the two molecules are no longer interlocked. At this point we have removed the entanglement of the two rings by something akin to a “sleight of hand”, but we have not yet solved the original problem, which was to build a configuration in which the two molecules are disentangled from each other in the 2D space. To achieve this, we must find a way to return the system from 3D to 2D while making sure that the two molecules remain disentangled during the transition.

**Figure 2.**
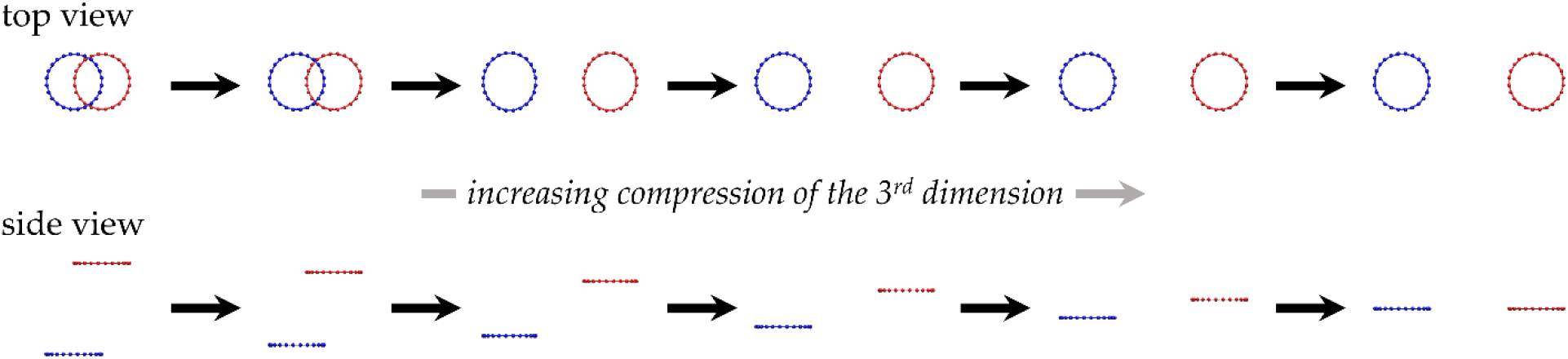
Snapshots from an energy minimization of the system shown in Figure 1 in which a temporary 3^rd^ spatial dimension is added and then gradually compressed out of existence. At the right hand side, the 3^rd^ dimension has been eliminated entirely and the two ring molecules have been returned to 2D.

We do this as follows. First, we introduce two half-harmonic potential functions to act as virtual “walls” that act only in the newly added 3^rd^ dimension and that prevent beads from assuming z-coordinates that exceed the values imposed by the walls. Next, we carry out an essentially conventional energy minimization during which the reference z-coordinates of the two virtual walls are gradually brought towards each other, until they eventually coincide. During this energy minimization, therefore, the spatial extent of the 3^rd^ dimension is gradually reduced until it becomes zero, at which point the two molecules have been effectively returned to 2D. Once the two virtual walls become coincident, we drop the z-coordinates of all beads (since by this point they should all be very similar to each other) and continue with a conventional energy minimization in 2D. Our hope is that as the 3^rd^ dimension becomes “squeezed” out of existence, the two molecules will begin to approach each other, and will be forced to accommodate each other by seeking to minimize their steric interactions. Assuming that this is done in a sufficiently gradual way, it should be possible for the molecules to find a non-clashing and disentangled configuration in 2D.

The ideas sketched above have been implemented in a simple molecular mechanics code, that has then been used to carry out an energy minimization of the system shown in Figure 1 (see Methods). Snapshots taken from selected points during that energy minimization are illustrated in Figure 2. Viewed from a vantage point perpendicular to the x,y plane, the two molecules appear to become disentangled from each other almost magically. Viewed from a vantage point perpendicular to the added dimension, however, the basis of the “trick” becomes clear: the two molecules effectively slide past each other as the walls that act upon them are moved toward each other. Movies illustrating the entire trajectory from both top and side views are provided in Supporting Movies S1A and S1B, respectively.

### Disentangling ring molecules that are interlocked in 3D

The above situation represents one in which a problem in 2D space is resolved by temporarily adding a 3^rd^ dimension and then compressing it until it disappears. To show that the same concept allows a problem in 3D space to be resolved by temporarily adding a 4^th^ dimension, we consider the system shown in Figure 3. Here the same ring molecules used above are now interlocked in 3D. To resolve the entanglement in this case, each ring is assigned a displacement in a 4^th^ spatial dimension (which we call “w”), and half-harmonic potential functions act as walls in this dimension. As before, we perform a single energy minimization, gradually moving the reference positions of the two walls towards their final values at w = 0, at which point the system has been returned to 3D. Snapshots taken from selected points during the energy minimization are illustrated in Figure 4, and a movie illustrating the entire trajectory is provided in Supporting Movie S2.

**Figure 3.**
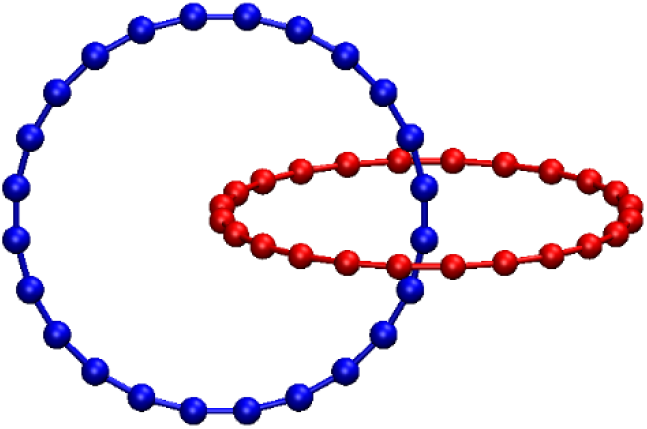
The same two ring molecules from Figure 1 but now interlocked in 3D.

**Figure 4.**
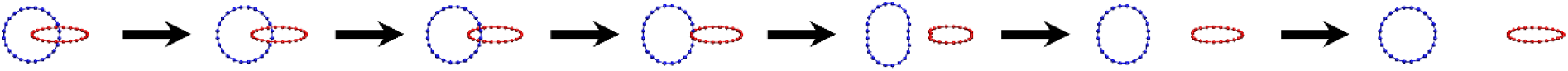
Snapshots from an energy minimization of the system shown in Figure 3 in which a temporary 4^th^ spatial dimension is added and then gradually compressed out of existence. At the right hand side, the 4^th^ dimension has been eliminated entirely and the two ring molecules have been returned to 3D.

A close examination of the behavior of the two rings during the trajectory illustrates some of the challenges in conceptualizing behavior that occurs in 4D but that can only be visualized as a projection on to 3D. At early stages of the simulation (i.e. when the two walls remain separated by a substantial distance in w), the rings undergo no structural deformations. As the walls begin to approach each other in the 4^th^ dimension, however, so do the two rings, and the presence of steric interactions between the two rings (which are not interlocked in the 4^th^ dimension!) causes both of them to compress inwards slightly at the position of their closest contact. As the walls continue to close in, however, the “crisis” passes: the two rings avoid each other in 3D, becoming disentangled, and the distortions temporarily induced in the circular structures begin to disappear as the bond lengths and angles strive to return to their equilibrium values.

### Removal of egregious steric interactions in superimposed molecules in 3D

The same principle outlined above can be used to resolve atrocious steric clashes of the kind that might be introduced in large-scale models that require multiple macromolecules to occupy the same region of space (e.g. as might be encountered when building models of intracellular environments). To provide a particularly challenging test case for illustrative purposes, ten identical copies of a coarse-grained (CG) protein model were directly superimposed upon each other at a single location in 3D space. Users of conventional molecular dynamics codes will know that such a situation will either cause an immediate crash (owing to many of the inter-bead distances being zero) or will cause drastic deformations in the structures as the molecules attempt to relieve their steric clashes. But with the molecules each initially displaced in the 4^th^ dimension such that their steric clashes are removed (when measured in 4D), the same energy minimization protocol used in the previous test cases ensures that all steric clashes are eventually relieved in 3D without distorting any of the molecules. Snapshots taken from selected points during the energy minimization are illustrated in Figure 5, and a movie illustrating the entire trajectory is provided in Supporting Movie S3. Interestingly, while a few of the protein molecules appear to “bud off” from each other during the early stage of the minimization, most of the activity occurs late on as the 4^th^ dimension is finally “squeezed” out of existence.

**Figure 5.**
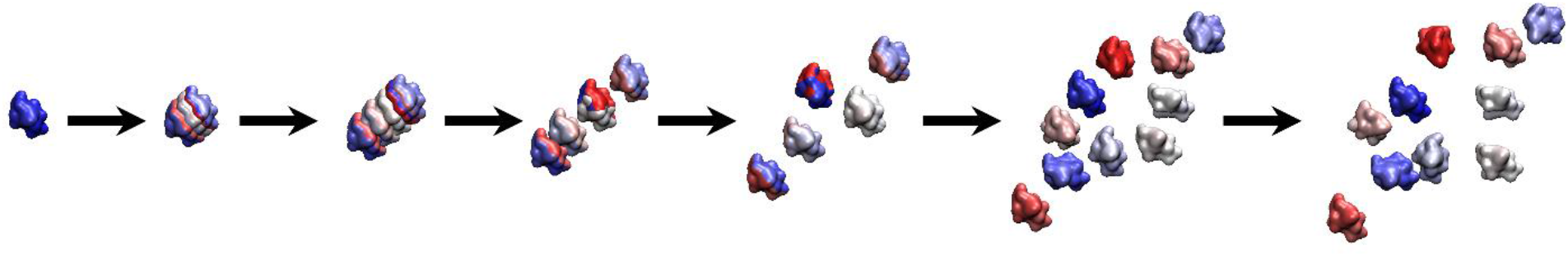
Snapshots from an energy minimization of ten superimposed coarse-grained protein models in which a temporary 4^th^ spatial dimension is added and then gradually compressed out of existence. At the right hand side, the 4^th^ dimension has been eliminated entirely and the ten protein models have been returned to 3D.

## Discussion

In this work I have shown that the temporary addition of a 4^th^ spatial dimension allows entanglements and steric clashes to be resolved in a largely automated fashion using an otherwise conventional molecular mechanics-based energy minimization. I have implemented the method in simple Fortran code (see Code availability), but for large-scale systems it is likely to be of more use when implemented within more popular simulation packages such as GROMACS [9], or OpenMM [10]. Making the necessary changes to existing simulation codes is, in principle, straightforward since all that is required is the addition of: (a) a 4^th^ spatial coordinate, and (b) adjustable half-harmonic potential functions that operate in this 4^th^ dimension. In practice, of course, the ease of implementing these changes will depend on the complexity of the pre-existing code and the extent to which it is documented.

One obvious area of application for the method presented here is to resolve the entanglements that can occur within double-helical regions of predicted RNA models [3] and the intertwining of DNA plectonemes that can, in principle, occur also in large-scale models of chromosomes. Since, however, similar entanglements can also occur in crudely constructed models of any macromolecule that contains ring structures (e.g. the sugar rings of carbohydrates or the aromatic sidechains of proteins), the general approach is likely to be applicable to a variety of macromolecular systems. A second area of application for the method is in resolving atrocious steric clashes that can be introduced when molecules are inadvertently superimposed upon each other during system construction. While there are, of course, other ways to solve such problems (see Introduction), the present method has the advantage of being both automatable and effectively agnostic to the shapes of the molecules. Moreover, it allows even highly overlapping and/or highly extended molecules to be easily separated from one another by displacing them by only a matter of a few Ångstroms in the added 4^th^ dimension.

Although the method has little in the way of adjustable parameters, the user is responsible for devising a way to distinguish and label those molecular entities that the user wishes to become, or to remain, disentangled or non-overlapping. In the case of systems that involve freely diffusing macromolecules, i.e. where entanglements and clashes are intermolecular in origin, this is easy to do. In more complicated cases, where entanglements or clashes might be intramolecular in nature (e.g. large RNAs or DNAs), it might make most sense to assign a different label to each double-helical region, single-stranded loop, or plectoneme. Once these labels have been assigned, all that is required is that structures that have been assigned different labels be given initial displacements in the 4^th^ dimension that ensure they are well separated from each other when their distance is measured (in 4D). Some care is also likely to be required in choosing the rate at which the reference positions of the 4^th^ dimensional half-harmonic potential functions are brought toward each other: if this is carried out too quickly, then particles may not have sufficient time to reach an accommodation with each other as they are forcibly returned to 3D. One way to improve behavior in this regard might be to replace the use of a simple energy minimization algorithm with a molecular dynamics or stochastic dynamics algorithm, suitably adjusted for 4D.

Finally, I note that, having taken the step of temporarily adding a 4^th^ spatial dimension, there seems little reason why one could not add further spatial dimensions. This might be especially advantageous in cases where many molecules simultaneously clash in the same region of 3D space. When only a single added dimension is allowed, molecules might need to be displaced by tens of Ångstroms in the 4^th^ dimension in order for all of the steric clashes that would otherwise be present in 3D to be initially resolved. With more added spatial dimensions, it ought to be possible to achieve mutual separation of all clashing molecules using combinations of more modest displacements in the added dimensions.

## Acknowledgments

This research was supported by a grant from the National Institutes of Health (R35 GM122466) to AHE and supported in part through computational resources provided by The University of Iowa.

## Author Contributions

Adrian H. Elcock: Conceptualization, Methodology, Software, Investigation, Writing – original draft preparation, Writing – review and editing, Supervision, Project Administration, Funding acquisition.

